# Secondary metabolites from food-derived yeasts inhibit virulence of *Candida albicans*

**DOI:** 10.1101/2020.08.14.251447

**Authors:** Lohith Kunyeit, Nawneet K Kurrey, Anu-Appaiah K A, Reeta P Rao

## Abstract

Beneficial microbes in the intestine are thought to control pathogen overgrowth by competing for limited nutrients. Our findings modify this prevailing paradigm of a passive, microbial antagonistic mode of action to an active, directed mechanism mediated by specific secondary metabolites. We describe two food-derived yeasts, *Saccharomyces cerevisiae* and *Issatchenkia occidentalis*, that inhibit virulence traits of *Candida albicans*, including hyphal morphogenesis, biofilms formation and adhesion to intestinal epithelial cells. These yeasts also protect the model host *Caenorhabditis elegans* from *C. albicans* infection. We demonstrate that the protective activity is primarily retained in the secretome of the beneficial yeasts and the protection they provide as a physical barrier is minimal. Mutant analysis demonstrates that phenylethanol and tryptophol are necessary for protection and experiments with commercially procured compounds indicates that they are sufficient to inhibit *C. albicans* virulence. We propose food-derived yeasts as an alternative or combination therapy to conventional antifungal therapy for *C. albicans* infection.

## Introduction

The polymorphic yeast, *Candida albicans* has been associated with a range of infection outcomes from superficial ones such as thrush and vaginitis to life-threatening invasive blood-stream infections. A multilateral interaction with the host immune system and host microbiota maintains this opportunistic pathogen in a commensal form in the gastrointestinal and urogenital tracts (Neville et al., 2015). However, a condition of immune suppression or disruption of the host-microbiota can cause severe mortality and morbidity (Calderone and Fonzi, 2001). As a part of our microbiome, *C. albicans* has been shown to have a complex interaction with other microbes. Yeast to hyphal transition plays a major role in modulating pathogenic interactions (Nobile and Mitchell, 2005, Jacobsen et al., 2012).

Secondary metabolites serve as environmental sensors that may respond to cell density and regulate key determinants of virulence such as biofilm development. For example, anaerobic conditions and nutrient deprivation regulates the production of aromatic alcohols in yeast (Ghosh et al., 2008, Chen and Fink, 2006). In the context of mixed microbial communities, secondary metabolites produced by one species have been shown to modulate virulence traits in other microbes (Chen and Fink, 2006). For example, *Pseudomonas aeruginosa* produces phenazine, and 3-oxo-C12 homoserine lactone which deters hyphal growth in *C. albicans* (Hogan et al., 2004; Hogan and Kolter, 2002). Furthermore, *C. albicans* shares a synergistic relationship with *Staphylococcus aureus* or *Streptococcus gordonii* and increases biofilm biomass on implanted medical devices and host surfaces, as well as evading the immune system (Lohse et al., 2018, Berman and Sudbery, 2002, Peters et al., 2010).

There are a limited number of drugs used to control *Candida* infection, most of which exhibit unpleasant side effects. Frequent usage of antifungal drugs drives resistance due to mutations, and overexpression of drug efflux pumps, and has posed a major clinical challenge (Ford et al., 2015). Furthermore, the complex structure of biofilms, which harbor a community of microorganisms facilitates resistance to antifungals (Lohse et al., 2018, Berman and Sudbery, 2002). These challenges warrant alternative solutions for *Candida* infections.

Beneficial yeasts such as *Saccharomyces cerevisiae* var. *boulardii* and *S. cerevisiae* have been shown to prevent the colonization and virulence of *C. albicans* (Pericolini et al., 2017, Murzyn et al., 2010, Jawhara and Poulain, 2007). We and others have previously reported that secreted factors from food-derived beneficial yeasts inhibited virulence traits of many pathogenic fungi (Kunyeit et al., 2019, Banjara et al., 2016). These food-derived yeasts have been considered beneficial in many cultural settings, yet their molecular mechanism of action has not been identified. Here we report the identification of two aromatic alcohols, tryptophol and phenylethanol which inhibit adhesion of *C. albicans in vitro, ex vivo* and were also active against mature biofilms. Furthermore, we used the model host *C. elegans* to demonstrate that these secondary metabolites are both necessary and sufficient to protect the host from *C. albicans* infection. Purified forms of these aromatic alcohols inhibited *C. albicans* hyphal morphogenesis and biofilm formation at ≥100μM concentrations and protected *C. elegans* from *C. albicans* infection. Because the rise of drug-resistant fungal infections undermines the limited repertoire of treatment options, we propose that beneficial yeasts could be used to control and prevent *Candida* infection.

## Results

### Food derived yeasts inhibit virulence traits of *Candida albicans*

Adhesion and filamentation are the primary determinants for *C. albicans* virulence. *C. albicans* must first adhere to abiotic surfaces then filament to form a three-dimensional biofilm. Beneficial yeast cell number 10^8^/ ml was empirically determined as effective against adhesion of *C. albicans* SC5314 (Kunyeit et al., 2019). To test the effect of the two food derived yeasts *Saccharomyces cerevisiae* and *Issatchenkia occidentalis* (Archana and Anu-Appaiah, 2018) on *C. albicans* virulence we tested adhesion, filamentation and biofilm formation *in vitro*. Adhesion to abiotic surfaces was tested under three different conditions detailed in the methodology section. Surfaces were treated with the beneficial yeast prior to inoculating *C. albicans* (pre-inoculation condition) or were inoculated at the same time (co-inoculation condition) as *C. albicans.* We also tested conditions where *C. albicans* was allowed to attach to the abiotic surface before exposure to beneficial yeast (post-inoculation condition). Our results indicate that the food-derived beneficial yeast, *S. cerevisiae* and *I. occidentalis*, inhibit adhesion of *C. albicans* to abiotic surfaces in all conditions tested (Figure 1A & B, figure supplement 1). Furthermore, the extent of inhibition observed with the beneficial yeast was comparable to the commercially available beneficial yeast *S. boulardii*, which has showed ∼85%, ∼60% and ∼38% in pre-inoculation, co-inoculation and post-inoculation condition respectively.

**Figure 1.**
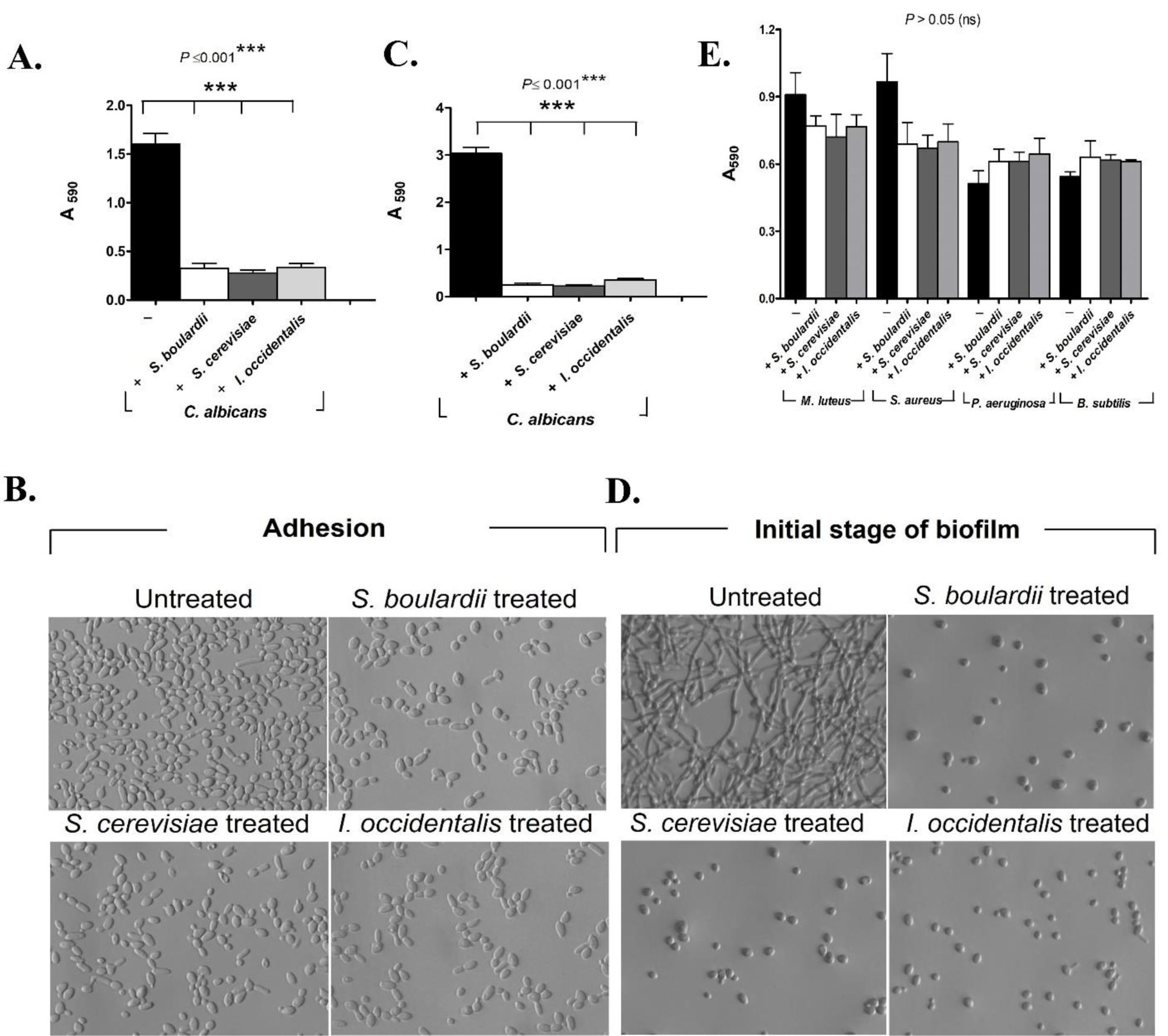
The food-derived, beneficial yeasts *S. cerevisiae, I. occidentalis* inhibit adhesion and biofilm formation of the opportunistic fungal pathogen, *C. albicans* but not bacterial species. *S. boulardii* was used as a reference control for the purpose of comparison. (A) Each yeast strain was co-incubated (represented as ‘+’) with *C. albicans* in RPMI and/or SC media. Staining with 0.5% crystal violet was used to quantify the adhesion of *C. albicans* on plastic surface (n = 5 experimental replicates). (B) A representative image showing *C. albicans* that remain attached to an abiotic surface when treated with beneficial yeasts. (C) *C. albicans* was treated with the beneficial yeasts for 24 h to test their effect on biofilm formation (n=8) and (D) microscopic illustration upon treatment with novel yeasts as compared to the reference strain. (E) Bacterial biofilms were treated with the beneficial yeasts, *S. cerevisiae* and *I. occidentalis* (n= 3). Error bars represents means± standard deviations (SD) and all values were represented at a significance with respect to control using One-Way ANOVA followed by *post hoc* analysis using Tukey’s *t* test.

Filamentation of *C. albicans* is an important virulence factor for deep tissue invasion, escape from immune cells and biofilm formation (Wilson et al., 2016). Beneficial yeasts *S. cerevisiae* and *I. occidentalis* inhibit filamentation of *C. albicans* (Figure 1C & D) to the same extent as commercially available beneficial yeast strain, *S. boulardii*. We investigated the effects of beneficial yeasts on *C. albicans* biofilm, in three distinct stages of biofilm development, initial (2-10 h), intermediate (24 h) and mature (48 and 72 h). Beneficial yeasts were able to inhibit biofilm formation significantly (∼79 %, *p*< 0.005, Figure 1C & D) on preformed *C. albicans* biofilms (up to 5-6 h old biofilms, under post-inoculation conditions) (Figure supplement 2A & B). However, intermediate (24 h) and mature biofilm (48 h) did not show any reduction in biomass but exhibited a marked decrease in metabolic activity (Figure supplement 3A & B) indicating decreased virulence. This result corroborates previous studies that show a positive correlation between higher metabolic activity of *Candida* cells in a biofilm with virulence (Cocuaud et al., 2005).

To test the effect of these novel yeasts on bacterial biofilms, we tested *Micrococcus luteus, Staphylococcus aureus, Pseudomonas aeruginosa* and *Bacillus subtilis*. We found that beneficial yeast treatment did not inhibit bacterial biofilm (Figure 1E) suggesting that these yeasts are not general microbial antagonists but rather exhibit specific antifungal activity against *C. albicans* and non-albicans *Candida* species (Kunyeit et al., 2019).

### Beneficial yeasts prevent attachment and invasion of *C. albicans* to human epithelial cells

We used Caco-2 cell lines, derived from human colon, to assess the ability of the beneficial yeasts to protect the epithelial cell from attachment and invasion of *C. albicans*. We measured inhibition under three experimental conditions: pre-inoculation; co-inoculation and post-inoculation, that mimic *in vitro* conditions tested. We assayed the ability of the putative beneficial yeasts, *S. cerevisiae* and *I. occidentalis*, to inhibit adhesion of *C. albicans* to live epithelial cells as compared to untreated samples, those that were not exposed to beneficial yeasts. We used the commercially available beneficial yeast *S. boulardii*, as a reference strain. Our results indicate that the beneficial yeasts were able to inhibit adhesion of *C. albicans* to cultured epithelial cells *ex vivo* under all conditions tested (Figure 2A & figure supplement 4). While pre-inoculation conditions practically abolished adhesion of *C. albicans* to Caco-2 monolayers, co-inoculation and post-inoculation condition significantly inhibited adhesion (94% and 84% respectively, *P*< 0.005). Furthermore, beneficial yeast treatment inhibited the morphological transition of *C. albicans* on Caco-2 cell monolayer (Figure 2B). Together these results suggest that presence of beneficial yeasts prevents attachment of *C. albicans* to both abiotic surfaces as well as live cells. Beneficial yeasts also inhibit filamentation of *C. albicans* which likely prevents biofilm formation on abiotic surfaces and invasion of live epithelial cells. To test whether beneficial yeasts are able to inhibit adhesion and filamentation of *C. albicans* in the presence of gastric juices, we simulated the gut environment *in vitro* using gastrointestinal components including pepsin, bile, pancreatic enzymes, electrolytes, ionic constituents and carbohydrates with pH of 2.5 (gastric juice) and 8.5 (bile juices) (Zhao et al., 2014). We found that beneficial yeasts were able to inhibit adhesion and germ tube development when applied 2 hours after *C. albicans* had attached to the abiotic surfaces (Figure supplement 5).

**Figure 2.**
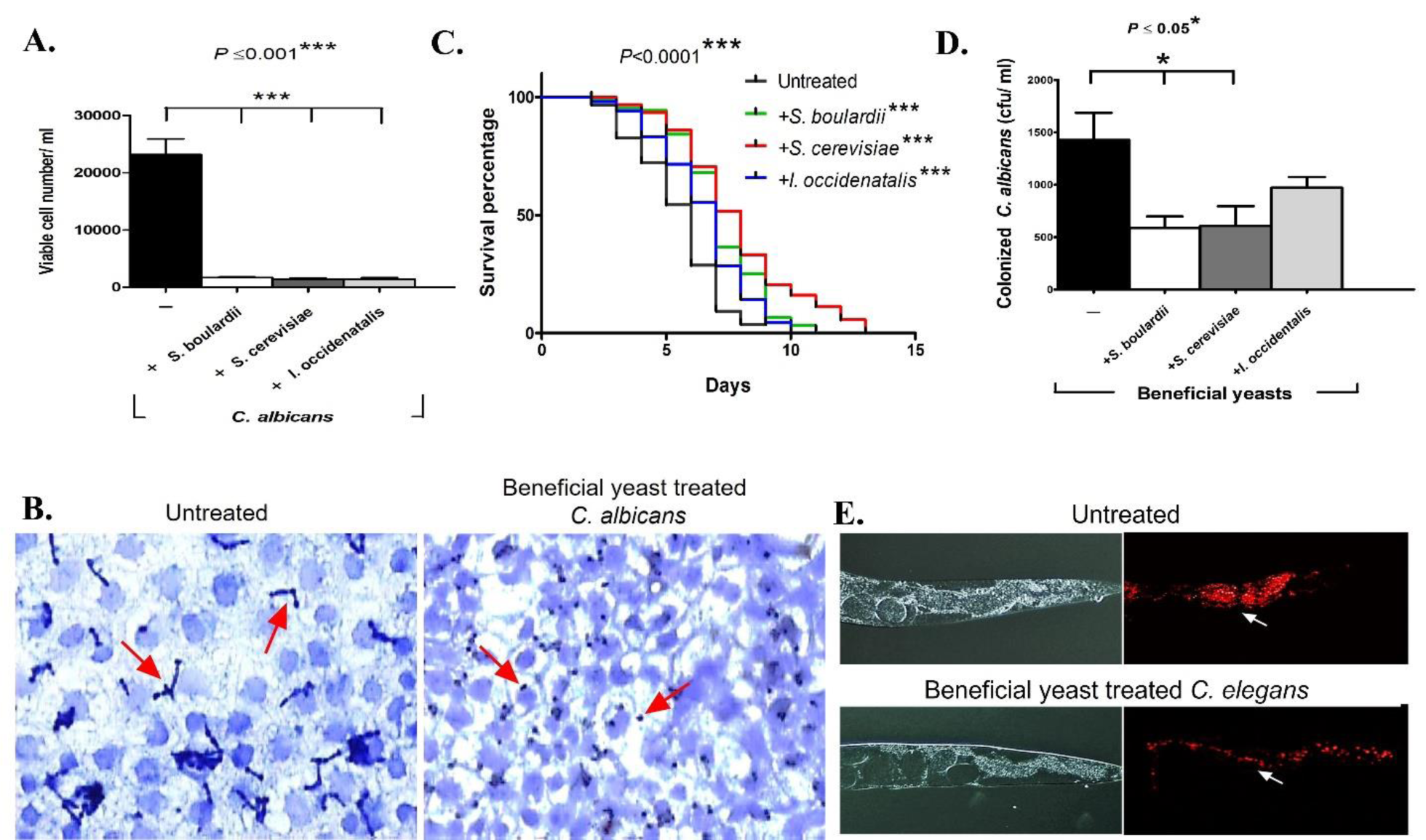
The beneficial yeasts prevent adhesion and invasion of epithelial cells and protect a live animal from *C. albicans* infection and colonization. (A) The novel yeasts *S. cerevisiae, I. occidentalis* were co-inoculated with *C. albicans* on Caco-2 monolayer. Adhesion of *C. albicans* to the epithelial layer was quantified by assessing colony forming units (CFU) on candida chrome agar (n=3 assay replicates). (B) Microscopic images of Caco-2 monolayers treated with beneficial yeasts (bottom) show *C. albicans* in ovoid yeast from cells as compared to untreated controls (top) where *C. albicans* germ tubes are visible indicating a transition to hyphal growth. (C) *C. elegans* was used as a live animal host. The life span of nematodes reared on a mixed diet of *C. albicans* and beneficial yeasts was plotted and compared to those reared on a diet of *C. albicans* alone (n = 4 experimental replicates, 143 ± 15 nematodes). (D) Co-exposure to beneficial yeasts reduced *C. albicans* colonization of the nematode gut. Colonization was quantified by counting colonies formed on candida chrome agar. Values were expressed as CFU/ ml (n=3) (E) as well as microscopically monitored using a mCherry tagged *C. albicans*. Error bars represent means± standard deviations (SD). Kaplan-Meier statistical analysis tools by Log-rank (Mantel-Cox) tests were used for the *C. elegans* survival assay and statistical significance is expressed with respect to control.

### Beneficial yeasts protect nematodes from *C. albicans* infections

*C. elegans* is an ideal experimental host system to study microbial interactions in the gastrointestinal tract. The nematode gut faithfully recapitulates a mammalian intestine in anatomy, innate immunity, and neuronal circuits (Elkabti et al., 2018, Schulenburg et al., 2004). Microbes introduced via the diet are established as the gut microbiome of the nematode thereby allowing the investigator to manipulate the intestinal microbiome. We leveraged these unique characteristics of *C. elegans* to study how the novel beneficial yeasts, *S. cerevisiae* and *I. occidentalis* interact with the pathogenic fungus, *C. albicans*, in the context of a live animal intestine. To mimic the experimental conditions used for *in vitro* and *ex vivo* studies we tested three exposure conditions. First, where worms were pre-exposed to beneficial yeasts prior to infection with *C. albicans* (pre-exposure), next, where worms were exposed to beneficial yeasts and *C. albicans* simultaneously (co-exposure) and finally, where *C. elegans* were treated with beneficial yeasts after infection with *C. albicans* (post-infection). Our results indicate that *C. elegans* that were exposed to beneficial yeasts were better able to cope with a *C. albicans* infection under all conditions tested (Figure 2C, figure supplement 6A & B). Worms that were exposed to *C. albicans* and beneficial yeast at the same time (co-exposure) extended host survival (by ∼3 days) as compared to the infected control group. We observed that, beneficial yeasts of the *Saccharomyces* species, *S. boulardii* or *S. cerevisiae* lived significantly longer than those treated with *I. occidentalis* in the co-exposure condition (Figure 2C). Worms that were exposed to beneficial yeasts prior to infection with *C. albicans* (pre-exposure) or treated with beneficial yeasts three days after infection (post-infection) with *C. albicans*, survived longer compared to untreated worms (Figure supplement 6A & B). Together these results indicate that application of beneficial yeasts at any point during the course of infection confers protective function and *C. albicans* infected worms are able to recover when beneficial yeasts are applied after *C. albicans* has presumably colonized the intestine. Therefore, we tested the hypothesis, that beneficial yeasts are able to inhibit colonization of worm gut with *C. albicans.* We used mCherry tagged *C. albicans* strain (Brothers et al., 2011) for microscopic observation of gut colonization and coupled that with colony forming units on differential growth medium, *Candida* chrome agar. Out results indicate that intestinal colonization is reduced upon beneficial yeasts treatment. Co-exposure of *S. cerevisiae* or *S. boulardii* significantly reduced colonization of *C. albicans* as compared to untreated *C. elegans.* Treatment with the non-*Saccharomyces* beneficial yeast, *I. occidentalis*, also reduced gut colonization but to a lesser extent (Figure 2D & E). The post-infection condition abolished intestinal colonization of *C. albicans* by day 4 of beneficial yeast treatment. Pre-exposure reduced *C. albicans* colonization during first two days, but in subsequent days the pathogen successfully recolonized. Together these studies suggest that beneficial yeasts alleviate colonization of the gut with *C. albicans* and beneficial yeast treatment is most effective when it is present during an active infection although it can provide limited protection prior to an infection.

### Secretome of beneficial yeast confer protective activity

To understand the mechanism of beneficial yeasts, we hypothesized that the beneficial yeasts either pose a physical barrier by occluding the intestinal epithelium or secrete small secondary metabolites that mediate beneficial yeast activity. To test whether the protective activity was retained in the secretome, we exposed *C. albicans* to the secretome of *S. cerevisiae, I. occidentalis* and *S. boulardii* using a two-chamber cell-culture insert (Figure 3A, figure supplement 7A). Individual live and heat-killed beneficial yeast strains were maintained in the upper chamber and *C. albicans* in the lower chamber of cell-culture insert wells so that cell-to-cell contact is prevented but small molecules present in the beneficial yeast secretome are able to diffuse to the lower chamber. Next, we tested whether these food-derived beneficial yeasts pose a physical barrier that inhibit adhesion of *C. albicans*. We introduced metabolically inactive, heat-killed beneficial yeast along with *C. albicans* in 96-well plates. We observed that, the heat-killed beneficial yeast partially inhibit adhesion due to physical barrier effect but *C. albicans* are able to filament and develop biofilms (Figure supplement 7 C, D & E). We further confirmed this finding using a contactless dual-chamber cell-culture experimental set up, where the secretome of heat-killed beneficial yeasts placed in upper chamber is unable to inhibit the virulence of *C. albicans* (Figure supplement 7B). This is in contrast to live beneficial yeasts which showed a robust inhibition of adhesion (Figure 3B) and hyphal development (Figure 3C) of *C. albicans*. Together these results demonstrate that the protective activity of beneficial yeasts is primarily retained in the secretome and the barrier function offers minimal protection.

**Figure 3.**
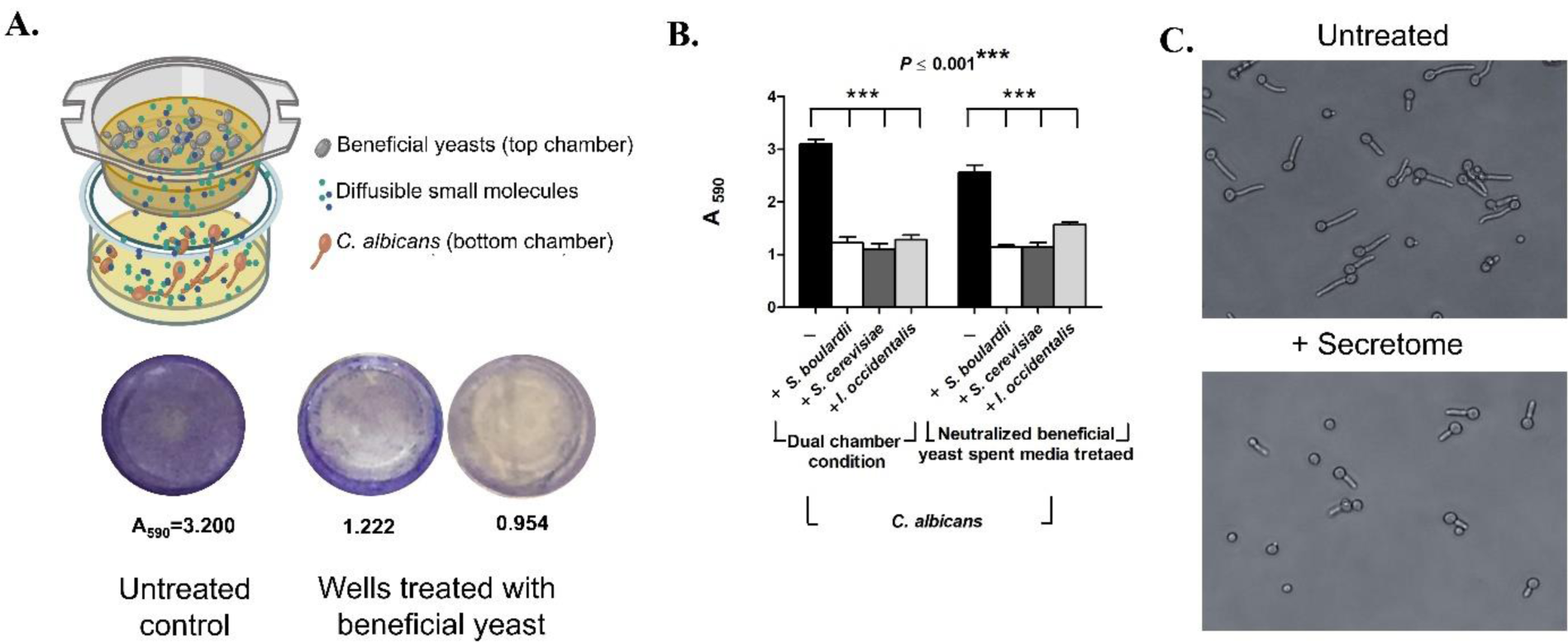
Diffusible metabolites from beneficial yeasts inhibit filamentation of *C. albicans*. (A) Two-chamber cell insert wells were used where, yeasts were inoculated in the upper chamber and *C. albicans* in lower chamber in SC media. Only small molecules are able to diffuse between the two chambers. After incubation lower *C. albicans* inoculated chamber was washed three times with PBS and stained with 0.5 % crystal violet. (B) Secretome that was dual chamber condition (left bars) or neutralized (right bars) from beneficial yeasts was exposed to *C. albicans* for 24 h *C. albicans* biomass was quantified using crystal violet staining (n=4 assay replicates). (C) Representative micrograph of *C. albicans* germ tube development when treated with the secretome from beneficial yeasts.

To confirm that this inhibitory effect was due to a secreted small molecule and not acidification of growth media that is known to inhibit hyphal development in *C. albicans* we neutralized, the spent media derived from *S. cerevisiae, I. occidentalis* and *S. boulardii* and tested the ability of this conditioned media to inhibit filamentation of *C. albicans*. Our results indicate that neutralized spent media also inhibited adhesion and filamentation of *C. albicans* (Figure 3B). Together these results suggest that secondary metabolites secreted by the beneficial yeasts, and not the acidic nature of the spent media, inhibited adhesion and filamentation of *C. albicans*. Simultaneously, we tested whether the secretome of beneficial yeasts was able to kill *C. albicans*. We used agar diffusion assay and found that exposure to the scretome of beneficial yeast did not reduce viability of *C. albicans*.

We and others have observed that beneficial yeast action is dose dependent, where cell number (10^8^/ ml) plays a key role in adhesion and hyphal inhibitory ability of *Candida* species (Lohith and Anu-Appaiah, 2018, Kunyeit et al., 2019). Furthermore, heat-killed beneficial yeasts were largely non-functional (Figure supplement 7). Therefore, we tested the hypothesis that secondary metabolites, responsive to cell density, inhibit virulence of *C. albicans*. We used Liquid Chromatography followed by tandem Mass Spectroscopy to LC-MS/ MS to separate and analyze the mass of the predicted secondary metabolites in the spent media. We confirmed the presence of two aromatic alcohols, tryptophol and phenylethanol, secreted by several yeast species (Chen and Fink, 2006, van Rijswijck et al., 2015) that have recently been shown to inhibit *C. albicans* filamentation (Chen and Fink, 2006, Martins et al., 2007). HPLC quantification revealed that the secretome of the beneficial yeasts *S. cerevisiae, I. occidentalis* and *S. boulardii* contained 140-150 μM of phenylethanol and 120-178 μM of tryptophol (Table 1, figure supplement 8). Next, we tested whether pure tryptophol or phenylethanol was able to inhibit virulence of *C. albicans* and confer host protection of *C. elegans* against *C. albicans* infection. We tested commercially available purified tryptophol and phenylethanol and confirmed that these compounds inhibit filamentation of *C. albicans* (Figure 4C). We found that tryptophol and phenylethanol effectively inhibited plastic adhesion (Figure 4A) as well as protected *C. elegans* from *C. albicans* infection (Figure 4B). Together these results indicate that the secondary metabolites tryptophol and phenylethanol are sufficient for the beneficial activity against *C. albicans*.

**Table 1.** Aromatic alcohol concentration of yeasts used in the study.

| Yeasts strains | Phenylethanol (μM) | Tryptophol (μM) |
| --- | --- | --- |
| *S. boulardii* | 154 ± 11.43 | 168 ± 06.27 |
| *S. cerevisiae* | 153 ± 14.18 | 120 ± 04.07 |
| *I. occidentalis* | 143 ± 07.36 | 178 ± 01.36 |
| *S. cerevisiae* (F1322) | 170 ± 18.03 | 164 ± 07.67 |
| *S. cerevisiae* (*aro8 aro9* mutants) | 51 ± 05.91 | 22 ± 04.89 |

**Figure 4.**
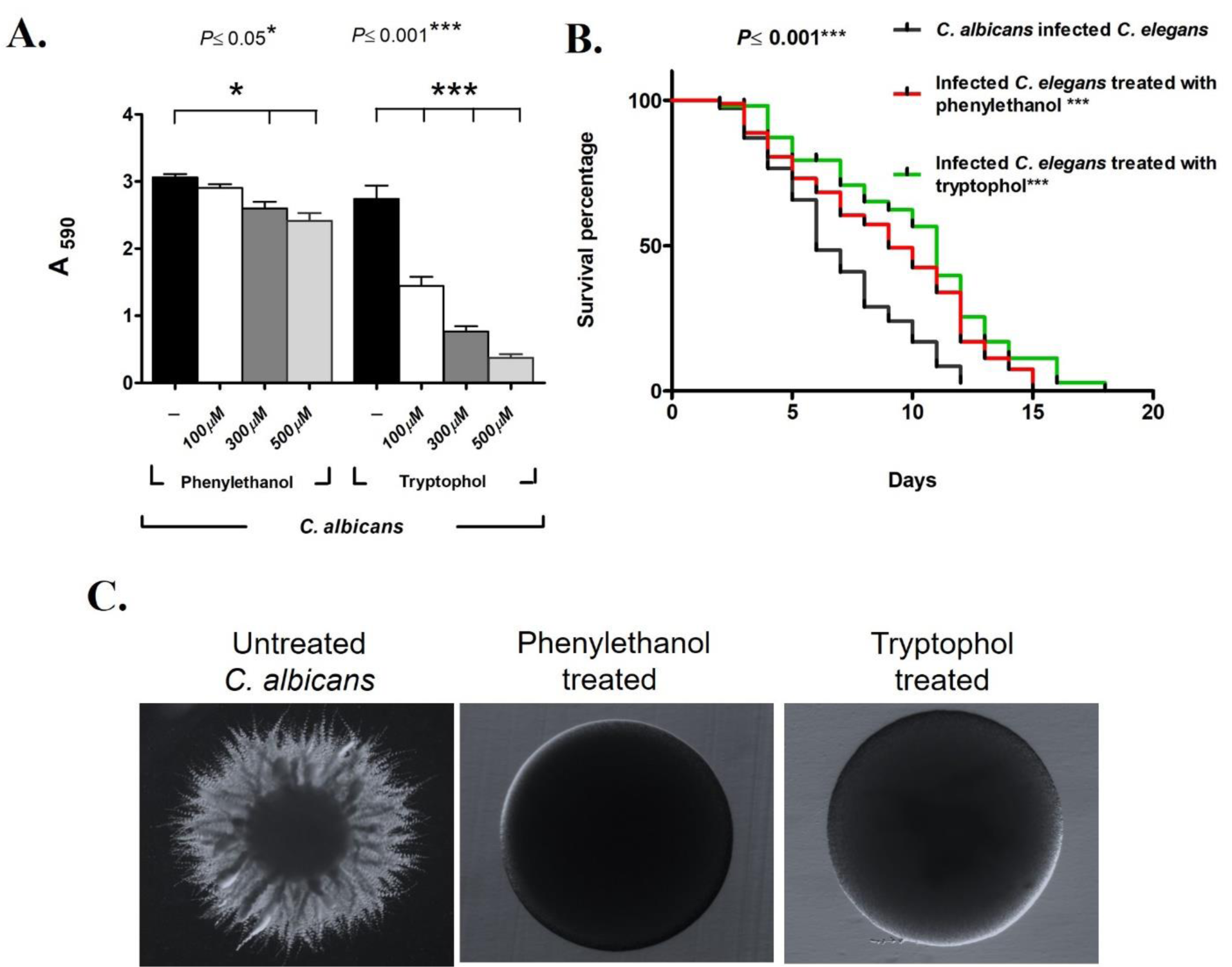
Phenylethanol and tryptophol are sufficient for beneficial activity *in vitro* and *in vivo.* (A) Varying concentration (100, 300 and 500 μM) of commercially procured aromatic alcohols, phenylethanol and tryptophol were exposed to *C. albicans* and biomass was measured with crystal violet (n = 5 experimental replicates). (B) Life span of *C. elegans* infected with *C. albicans* treated tryptophol (100 μM) (green curve) and phenylethanol (100 μM) (red curve) (n =3 experimental replicates, 82 ± 28 nematodes). (C) Representative images showing filamentation of *C. albicans*, upon phenylethanol and tryptophol treatment. Error bars represents means± standard deviations (SD). Kaplan-Meier statistical analysis tools by Log-rank (Mantel-Cox) tests were used for the *C. elegans* survival assay.

### Mutants that do not produce the secondary metabolites are ineffective

To test the hypothesis that yeast derived aromatic alcohols are necessary for protection against *C. albicans*, we compared a laboratory strain of *S. cerevisiae* (F1322, Table supplement 1), and the isogenic double mutant *aro8 aro9*, which is unable to produce these aromatic alcohol (Chen and Fink, 2006). We used neutralized spent media to test the ability the secretome from the *aro8 aro9* mutant to inhibit adhesion and filamentation of *C. albicans* compared to the isogenic wild-type control. Our results indicate that wild-type yeast, inhibits virulence of *C. albicans* as compared to the isogenic double mutant *aro8 aro9* that is unable to produce tryptophol and phenylethanol (Figure 5A). To confirm these results *in vivo*, we co-infected *C. elegans* with *C. albicans* and either the wild type control strain or the *aro8 aro9* mutant strain. Our result demonstrates that nematodes co-infected with wild type *S. cerevisiae* survived longer than those that were co-infected with the *aro8 aro9* mutant (Figure 5B) suggesting that *S. cerevisiae* produce aromatic alcohols that inhibit virulence traits of *C. albicans*, thereby protecting the nematode host from *C. albicans* infection. These laboratory strains were unable to thrive in the gastrointestinal juices as compared to the food derived yeasts, suggesting that they likely not suitable for therapeutic purposes. Together these provide a molecular mechanism for the action of food-derived beneficial yeasts where diffusible secondary metabolites, tryptophol and phenylethanol, contained in their secretome are largely responsible for their protective properties against the opportunistic pathogenic fungi *C. albicans* (Figure 5C). This report demonstrates that tryptophol and phenylethanol are necessary and sufficient for the beneficial activity of the novel food derived yeasts.

**Figure 5.**
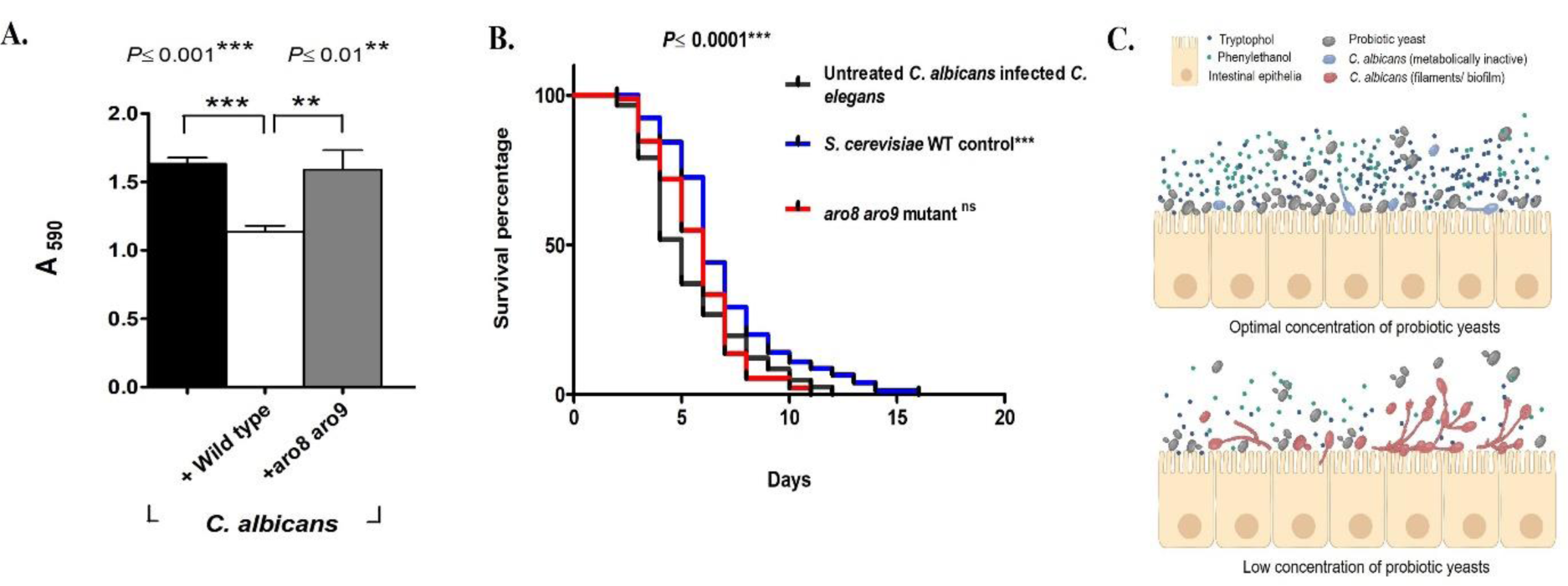
*S. cerevisiae* mutant analysis shows phenylethanol and tryptophol are required for beneficial activity *in vitro* and *in vivo.* (A) The neutralized secretome of *aro8 aro9* double mutant (grey bar), that does not produce aromatic alcohols, was tested for its ability to prevent adhesion of *C. albicans*, compared to its isogenic wild type counterpart (white bar) or untreated controls (black bars), (n=4 assay replicates). (B.) Life span of *C. elegans* treated either with *aro8 aro9* double mutant or its wild type counterpart. The double mutant (red curve) does not protect the nematode as well as its isogenic wild type control strain (blue curve) (n= 4 experimental replicates, 163 ± 11 nematodes). (C) Working model for the mechanism by which beneficial yeasts function to inhibit *C. albicans* virulence. Error bars represents means± standard deviations (SD). Kaplan-Meier statistical analysis tools by Log-rank (Mantel-Cox) tests were used for the *C. elegans* survival assay and statistical significance is expressed with respect to control.

## Discussion

Morphological transition of *C. albicans* is a major virulence factor (Nobile and Mitchell, 2005) that is associated with biofilm development on implanted medical devices and, host epithelium, and that helps the pathogen evade the immune system (Lohse et al., 2018, Berman and Sudbery, 2002). We show that food derived beneficial yeasts *S. cerevisiae* and *I. occidentalis*, are able to inhibit virulence traits of *C. albicans* and prevent infection in a live animal.

Mature biofilms are highly resistant to various therapeutic agents. Commercial antifungals such as fluconazole, caspofungin and amphotericin B reduce metabolic activity (Tobudic et al., 2010). While application of the putative beneficial yeasts did not decrease biofilm biomass of mature biofilms, they significantly reduced (*P*< 0.05) metabolic activity of the biofilm. Application of cell-free, neutralized spent media from beneficial yeasts, also reduced the metabolic activity of 24h old *C. albicans* biofilm by 26% suggesting that soluble metabolites play a crucial role in controlling *Candida* growth. It has been suggested that reduced viability rather than complete elimination of the biomass and filamentation may be a better avenue for keeping *Candida* in its commensal state. This decrease in metabolic activity has been proposed as a mechanism for beneficial effect of *Lactobacillus rhamnosus* (Ribeiro et al., 2017).

Yeasts produces aromatic alcohols such as tryptophol and phenylethanol as a cell density manner in the SC media of low ammonium sulphate (Chen and Fink, 2006). The yeasts cell number more than the 10^7^/ ml produce 140-200μM of tryptophol and phenylethanol (Avbelj et al., 2015) and our study showed that beneficial yeast cell number of 10^8^/ ml produce 120-170 μM and these concentrations of aromatic alcohols tryptophol and phenylethanol inhibit the *C. albicans* filamentation and reduces the biomass. Previous reports have shown that intraperitoneal administration of phenylethanol (70μM) and other *C. albicans* autoregulatory alcohols have protected mice from invasive Candidiasis (Martins et al., 2012). Similarly, pretreatment of tryptophol (200 μM) to *Galleria mellonella*, a greater wax moth significantly enhanced the host survival from the *Candida* infection (Singkum et al., 2019). The conditions of the gastrointestinal tract have been shown to be favorable for the production of aromatic alcohols (Ghosh et al., 2008). In line with these published reports, our observations in simulated gastrointestinal tract condition indicates that treatment with beneficial yeast inhibits *Candida* virulence. The gastrointestinal tract is a complex environment where the beneficial microbes are typically confined to the gut where they have a local effect. However, the diffusible aromatic alcohols they produce are able to mount a systemic and long ranging response.

Overall, this study demonstrates that food-derived beneficial yeasts produce aromatic alcohols, specifically tryptophol and phenylethanol, that inhibit virulence traits of *Candida albicans*. Since heat-inactivated beneficial yeasts retain minimal activity, we propose that beneficial yeasts present a chemical inhibition as well as pose a physical (passive) barrier that prevent *Candida* infections. Therefore, they may be used as an alternative to or in combination with traditional antifungal drugs.

## Materials and methods

### Microbial strains, media preparation and chemical used

Bacteria and yeasts strains were maintained in the Luria broth (LB) and Yeast extract Peptone Dextrose (YPD) respectively unless otherwise mentioned (table supplement 1). RPMI and/or synthetic complete (SC) media was used to study adhesion and biofilm experiments. Pepsin, ox-bile, pancreatin Dulbecco’s Modified Eagle Medium (DMEM) and fetal bovine serum (FBS) were procured from HiMedia Laboratories Pvt. Ltd. RPMI-1640, yeast synthetic drop-out medium supplements, yeast nitrogen base, tryptophol, phenylethanol and 3-(4,5-Dimethylthiazol-2-yl)-2,5-Diphenyltetrazolium Bromide (MTT) were obtained from Sigma-Aldrich.

Spider media was used to monitor filamentation of *C. albicans* on agar plates. Briefly, *C. albicans* was spotted on spider agar plate containing 300, 500 and 1000 μM of tryptophol and phenylethanol and incubated for 4 days at 30 °C. Images were captured on Zeiss dissection microscope equipped with SPOT imaging software. To monitor the effects of aromatic alcohols, phenylethanol and tryptophol on *C. albicans* in liquid media, *C. albicans* (10^6^/ ml) was inoculated in SC media containing a range of (100-500 μM) of phenylethanol or tryptophol and incubated at 37 °C for 8-10 h. Adhesion of *C. albicans* was quantified as previously described (Jin et al., 2003). Briefly, the plates were air-dried and incubated with 50 *μ*l of 0.5% crystal violet for 25 min. Adherent cells were washed with PBS, distained with 95% (v/v) ethanol and quantified by measuring absorbance at 590 nm.

To prepare secretomes, cultures were maintained for 24 h in Synthetic Complete (SC) media with 50 μM ammonium sulphate at 37 °C in with aeration. Cell pellets were removed by centrifugation. The pH of the remaining spent media was neutralized, depleted nutrients were replenished, and filter sterilized prior to testing.

Simulated gastric and bile juices were prepared as previously described (Casey et al., 2004). Gastric juice was prepared using glucose 3.5 g/ L, sodium chloride 2.05 g/ L, potassium phosphate monobasic 0.6 g/ L, calcium chloride 0.11 g/ L, potassium chloride 0.37 g/ L, pepsin 0.7 g/ L with pH adjusted to 2.5. Bile juice media contained a glucose 3.5 g/ L, sodium chloride 2.05 g/ L, potassium phosphate monobasic 0.6 g/ L, pancreatin 0.750 g/ L and bile 0.5 g/ L with pH 8.50. The reference strain *S. boulardii*, or novel yeast strains, *S. cerevisiae* and *I. occidentalis* (10^8^/ ml) were co-inoculated with *C. albicans* (10^7^/ ml) and incubated for 180 min in simulated gastric juice and bile juices. Adherent cells were quantified by crystal violet staining described above.

### *In vitro* plastic adhesion and biofilm assays

*In vitro* assays were performed in pre-inoculated, co-inoculated and post-inoculated conditions as perviously described (Kunyeit et al., 2019). For pre-inoculated experiments, the beneficial yeasts were first inoculated (10^8^/ ml) into 96-well plates and incubated for 30 min at 37 °C. *C. albicans* cells (10^7^/ ml) were then introduced and incubated for an additional 90 min. For co-inoculated conditions, the beneficial yeasts and *C. albicans* were incubated together for 120 min at 37 °C. For post-inoculated condition, *C. albicans* was inoculated to the 96-well plates for 30 min before beneficial yeast were introduced on the *C. albicans* cells that had adhered to the bottom of the well and incubated for another 90 min. After treatment, plates were washed three times with phosphate buffer saline (PBS, pH 7.4) to remove unattached cells. Adherent *C. albicans* cells were either microscopically visualized or quantified as described (Jin et al., 2003, Parolin et al., 2015). Pre-inoculation and co-inoculation conditions test the hypothesis that the beneficial yeasts are prophylactic while the post-inoculation condition tests whether beneficial yeasts maybe used as a therapeutic.

We also tested the effect of beneficial yeasts on the adhesion of *Staphylococcus aureus, Micrococcus luteus, Pseudomonas aeruginosa* and *Bacillus subtilis.* Bacterial culture (optical density of 0.5) and 10^8^/ ml of beneficial yeast was maintained in RPMI/SC media and incubated for 48 h at 37 °C. After incubation, unattached cells were removed and attached bacterial cells were quantified by 0.5% crystal violet (Jin et al., 2003).

*C. albicans* biofilms were tested as described earlier (Kunyeit et al., 2019) at three distinct biofilm development phases initial (0–10 h), intermediate (24 h), and mature (48 h) (Jabra-Rizk et al., 2004). Initial stages of biofilm were tested in two ways. Co-inoculated treatment, where *C. albicans* (10^6^/ ml) and beneficial yeasts (10^8^ /ml) were co-incubated for 24 h at 37 °C or post-inoculated condition, where *C. albicans* was allowed to attach and initiate biofilm formation for 90 min prior to introduction of 10^8^ /ml beneficial yeasts. Plates were incubated 24 h at 37 °C in mild shaking (90 rpm). Intermediate and mature biofilm treatments were conducted on 24 and 48 h old pre-formed *C. albicans* biofilm respectively. Briefly, 10^6^/ ml of *C. albicans* was inoculated in to RPMI/SC media for 30 min. To develop a stable biofilm, unadhered *C. albicans* cells were removed and incubated at 37 °C, for an additional 24 h or 48 h for intermediate or mature biofilms respectively. 10^8^/ ml of beneficial yeasts were then introduced to these biofilms and incubated for another 24 h. Crystal violet staining was used for quantifying the biofilm of *C. albicans* which explained above. The metabolic activity of biofilm was checked by MTT assay (Mosmann, 1983).

### *Ex vivo* adhesion of *C. albicans* to intestinal epithelium

Caco-2, the standard human epithelial colon cell lines were used to monitor the effect of beneficial yeasts on interaction between *C. albicans* and intestinal epithelia. 5×10^4^ cells/ mL of Caco-2 were seeded in to 96-well microtiter plate containing DMEM media with 10% FBS and incubated at 37 °C with 5% CO2. The experiment was performed as described earlier (Kunyeit et al., 2019). After 20 days, the resulting monolayer of Caco-2 was treated with *C. albicans* and beneficial yeasts. Experiments was conducted in pre-inoculated, co-inoculated and post-inoculated conditions analogous to the plastic adhesion experiment described above. *C. albicans* colony forming units (CFUs) were assessed using candida-chrome agar plate. Simultaneously, crystal violet stain (0.5%) was used to stain the adhered yeasts on Caco-2 cell monolayer and images were taken using bright field microscope.

### *In vivo C. elegans* infection and colonization assay

*C. elegans* was used as a model host to monitor the effects of beneficial yeast upon *C. albicans* infection (Jain et al., 2009). For this *in vivo* assay, nematode eggs were harvested from six to eight worms reared on Nematode growth media (NGM) agar plate containing *E. coli* OP50 at 20 °C for three days. 40-50 eggs were transferred to fresh NGM plate contained *E. coli* OP50. Assays were conducted in pre-exposure, co-cultured and post-exposure condition (Kunyeit et al., 2019). For pre-exposure, L3-L4 level worms were maintained on a diet of beneficial yeasts for two days and then transferred to the *C. albicans* diet. Viability was scored daily, and worms reared on the regular diet of *E. coli*, OP50 prior to infection with *C. albicans* was used as an untreated control group. For co-exposure worms were raised on mixed cultures of beneficial yeasts (10^6^ cells) and *C. albicans* (10^6^ cells) while for post-infection treatment, worms were first infected with *C. albicans* (10^6^ cells, for 2 days) and subsequently transferred the worms into beneficial yeast lawn (10^6^ cells / 20 μL). *C. albicans* infected worms that were treated with the beneficial yeast, were compared with the control where *C. albicans* infected worms were transferred in OP50 lawn.

Five to six worms grown in pre-exposures, co-exposure or/and post-infection condition were allowed to crawl on a fresh unseeded NGM plate to remove yeast or *E. coli* cells that might have attached to the outer cuticle of the nematode. Washed worms were transferred to an eppendorf tube containing 100 μl Phosphate Buffered Saline (PBS) then crushed using pellet pestle and vortexed for 3-4 min. Finally, viable colony forming units (CFUs) were counted of *C. albicans* by plating appropriate dilution on candida chrome agar to differentiate *C. albicans* cells from beneficial yeast isolates. To visualize colonization of the nematode gut with *C. albicans*, an mCherry tagged *C. albicans* was used to capture images using a Zeiss Axiovert 200M florescence Microscope.

### Preparation of heat-killed beneficial yeast cells

10^8^/ ml beneficial yeasts were incubated at 70-75°C for 30 min in the water bath. Then, washed three times with PBS buffer. Cell death was confirmed by using methylene blue staining.

### Two-chamber cell insert assay to test the secretome from beneficial yeasts

To assess the beneficial yeast metabolite effect on *C. albicans* a dual chamber cell-insert chamber was used. The dual chamber cell insert has a filter (0.4 μm pore size) which separate the beneficial yeasts from *C. albicans* cells. Briefly, 10^8^ cells /ml of beneficial yeasts (live and heat-killed cells in the indivdual experiment) and 10^6^ cells /ml of *C. albicans* strain was maitained in upper and lower chamber respectively and incubated for 24 h at 37 °C with mild shaking (90 rpm). After incubation, discarded the upper beneficial yeast contained chamber and lower part of the plate contained *C. albicans* was washed three times with phosphate buffer saline (PBS, pH 7.4) to remove unadhered *C. albicans* cells. Finally, quantified by 0.5% of crystal violet staining (Kunyeit et al., 2019).

### HPLC and LC-MS/ MS analysis of tryptophol and phenylethanol

Supernatant from 24 h old cultures of the two novel yeasts *S. cerevisiae* and *I. occidentalis* or the reference strain *S. boulardii* was used to quantify tryptophol and phenylethanol by High Pressure Liquid Chromatography (Shimagzu corporation, Japan). 10^8^/ ml yeasts cells were cultured in SC media contained low ammonium sulphate (50 μM) for 24 h at 37 °C. The supernatant was collected by centrifugation and filtered and used it for HPLC analysis using a C18 column. 0.01 % (vol/vol) trifluoroacetic acid (TFA) in water used as the first solvent system A followed by 0.01 % TFA in acetonitrile as the next solvent system B. The flow rate was maintained at 1.0 mL/ min for 12 min, with a 10 % of solvent B and increased to 25 % in 1.0 min and continued for 15 min; then raised to 100 % of solvent B in 1.0 min and maintained it for 3 min. Finally, solvent B was decreased to 10 % for 1 min and maintained it for 8 min and detected at 210 nm. To confirm these aromatic alcohols Hybrid Quadrupole-TOF LC-MS/ MS (Sciex Triple ToF 5600, Singapore) method was used with the solvent system which mentioned as above (Li et al., 2008).

### Statistical analysis

Statistical significance was assessed by One-way analysis of variance (ANOVA) followed by *post hoc* analysis using Tukey’s *t* test at a significance level of *P* < 0.05. Results were expressed as means± standard deviations (SD). Analyses were performed with GraphPad Prism 5 software (GraphPad Software Inc., San Diego, CA, USA). Kaplan-Meier statistical analysis tools were used for the *C. elegans* survival assay.

## Acknowledgements

We thank the Director, CSIR-Central Food Technological Research Institute (CFTRI) for encouragement and research support; LK is grateful to the INSPIRE program, Department of Science and Technology, Government of India and Fulbright-Nehru doctoral fellowship, United States-India Education Foundation (USIEF), India for the financial support for his doctoral research. This work is partially supported by NIH-NCCIH 1R15AT009926-01 grant to RPR. We thank Dr. Elizabeth F. Ryder, and Dr. José M. Argüello, Worcester Polytechnic Institute, MA, for their critical review of the manuscript and Prof. Gerald R. Fink, MIT, Cambridge, MA, for providing *S. cerevisiae* strains F1322 and L8184.

## References

Archana, K. M. & Anu-Appaiah, K. A. 2018. Detection system for *Saccharomyces cerevisiae* with phenyl acrylic acid decarboxylase gene (PAD1) and sulphur efflux gene (SSU1) by multiplex PCR. Archives of Microbiology, 200, 275–279.

Avbelj, M., Zupan, J., Kranjc, L. & Raspor, P. 2015. Quorum-sensing kinetics in *Saccharomyces cerevisiae*: A symphony of ARO genes and aromatic alcohols. Journal of Agricultural and Food Chemistry, 63, 8544–8550.

Banjara, N., Nickerson, K. W., Suhr, M. J. & Hallen-Adams, H. E. 2016. Killer toxin from several food-derived *Debaryomyces hansenii* strains effective against pathogenic *Candida* yeasts. International Journal of Food Microbiology, 222, 23–29.

Berman, J. & Sudbery, P. E. 2002. *Candida albicans*: a molecular revolution built on lessons from budding yeast. Nature Reviews Genetics, 3, 918–30.

Brothers, K. M., Newman, Z. R. & Wheeler, R. T. 2011. Live imaging of disseminated candidiasis in zebrafish reveals role of phagocyte oxidase in limiting filamentous growth. Eukaryotic Cell, 10, 932–44.

Calderone, R. A. & Fonzi, W. A. 2001. Virulence factors of *Candida albicans*. Trends in Microbiology, 9, 327–35.

Casey, P. G., Casey, G. D., Gardiner, G. E., Tangney, M., Stanton, C., Ross, R. P., Hill, C. & Fitzgerald, G. F. 2004. Isolation and characterization of anti-*Salmonella* lactic acid bacteria from the porcine gastrointestinal tract. Letters in Applied Microbiology, 39, 431–8.

Chen, H. & Fink, G. R. 2006. Feedback control of morphogenesis in fungi by aromatic alcohols. Genes and Devlopment, 20, 1150–61.

Cocuaud, C., Rodier, M. H., Daniault, G. & Imbert, C. 2005. Anti-metabolic activity of caspofungin against *Candida albicans* and *Candida parapsilosis* biofilms. Journal of Antimicrobial Chemotherapy, 56, 507–512.

Elkabti, A. B., Issi, L. & Rao, R. P. 2018. *Caenorhabditis elegans* as a model host to monitor the *Candida* infection processes. Journal of Fungi, 4, 123.

Ford, C. B., Funt, J. M., Abbey, D., Issi, L., Guiducci, C., Martinez, D. A., Delorey, T., Li, B. Y., White, T. C., Cuomo, C., Rao, R. P., Berman, J., Thompson, D. A. & Regev, A. 2015. The evolution of drug resistance in clinical isolates of *Candida albicans*. Elife, 4, e00662.

Ghosh, S., Kebaara, B. W., Atkin, A. L. & Nickerson, K. W. 2008. Regulation of aromatic alcohol production in *Candida albicans*. Appllied Environmental Microbiology, 74, 7211–8.

Hogan, D. A. & Kolter, R. 2002. *Pseudomonas-Candida* interactions: An ecological role for virulence factors. Science, 296, 2229–2232.

Hogan, D. A., Vik, A. & Kolter, R. 2004. A *Pseudomonas aeruginosa* quorum-sensing molecule influences *Candida albicans* morphology. Moleclar Microbiology, 54, 1212–23.

Jabra-Rizk, M. A., Falkler, W. A. & Meiller, T. F. 2004. Fungal biofilms and drug resistance. Emerging Infectious Diseases, 10, 14–9.

Jacobsen, I. D., Wilson, D., Wachtler, B., Brunke, S., Naglik, J. R. & Hube, B. 2012. *Candida albicans* dimorphism as a therapeutic target. Expert Review of Anti-Infective Therapy, 10, 85–93.

Jain, C., Yun, M., Politz, S. M. & Rao, R. P. 2009. A pathogenesis assay using *Saccharomyces cerevisiae* and *Caenorhabditis elegans* reveals novel roles for yeast AP-1, Yap1, and host dual oxidase BLI-3 in fungal pathogenesis. Eukaryotic Cell, 8, 1218–27.

Jawhara, S. & Poulain, D. 2007. *Saccharomyces boulardii* decreases inflammation and intestinal colonization by *Candida albicans* in a mouse model of chemically-induced colitis. Medical Mycology, 45, 691–700.

Jin, Y., Yip, H. K., Samaranayake, Y. H., Yau, J. Y. & Samaranayake, L. P. 2003. Biofilm-forming a ability of *Candida albicans* is unlikely to contribute to high levels of oral yeast carriage in cases of human immunodeficiency virus infection. Journal of Clinical Microbiology, 41, 2961–2967.

Kunyeit, L., Kurrey, N. K., Anu-Appaiah, K. A. & Rao, R. P. 2019. Probiotic yeasts inhibit virulence of Non-*albicans Candida* Species. mBio, 10.

Li, M., Yang, Z. X., Hao, J. G., Shan, L. J. & Dong, J. J. 2008. Determination of Tyrosol, 2-phenethyl alcohol, and tryptophol in beer by High-Performance Liquid Chromatography. Journal of the American Society of Brewing Chemists, 66, 245–249.

Lohith, K. & Anu-Appaiah, K. A. 2018. Antagonistic effect of *Saccharomyces cerevisiae* KTP and *Issatchenkia occidentalis* ApC on hyphal development and adhesion of *Candida albicans*. Medical Mycology, 56, 1023–1032.

Lohse, M. B., Gulati, M., Johnson, A. D. & Nobile, C. J. 2018. Development and regulation of single- and multi-species *Candida albicans* biofilms. Nature Reviews Microbiology, 16, 19–31.

Martins, M., Henriques, M., Azeredo, J., Rocha, S. M., Coimbra, M. A. & Oliveira, R. 2007. Morphogenesis control in *Candida albicans* and *Candida dubliniensis* through signaling molecules produced by planktonic and biofilm cells. Eukaryotic Cell, 6, 2429–2436.

Martins, M., Lazzell, A. L., Lopez-Ribot, J. L., Henriques, M. & Oliveira, R. 2012. Effect of exogenous administration of *Candida albicans* autoregulatory alcohols in a murine model of hematogenously disseminated candidiasis. Journal of Basic Microbiology, 52, 487–491.

Mayer, F. L., Wilson, D. & Hube, B. 2013. *Candida albicans* pathogenicity mechanisms. Virulence, 4, 119–128.

Mosmann, T. 1983. Rapid colorimetric assay for cellular growth and survival: application to proliferation and cytotoxicity assays. Journal of Immunological Methods, 65, 55–63.

Murzyn, A., Krasowska, A., Stefanowicz, P., Dziadkowiec, D. & Lukaszewicz, M. 2010. Capric acid secreted by *Saccharomyces boulardii* inhibits *Candida albicans* filamentous growth, adhesion and biofilm formation. Plos One, 5, e12050.

Neville, B. A., D’Enfert, C. & Bougnoux, M. E. 2015. *Candida albicans* commensalism in the gastrointestinal tract. FEMS Yeast Research, 15, 1–13.

Nobile, C. J. & Mitchell, A. P. 2005. Regulation of cell-surface genes and biofilm formation by the *Candida albicans* transcription factor Bcr1p. Current Biology, 15, 1150–5.

Parolin, C., Marangoni, A., Laghi, L., Foschi, C., Nahui Palomino, R. A., Calonghi, N., Cevenini, R. & Vitali, B. 2015. Isolation of vaginal lactobacilli and characterization of anti-*Candida* activity. PLoS One, 10, e0131220.

Pericolini, E., Gabrielli, E., Ballet, N., Sabbatini, S., Roselletti, E., Cayzeele Decherf, A., Pelerin, F., Luciano, E., Perito, S., Justen, P. & Vecchiarelli, A. 2017. Therapeutic activity of a *Saccharomyces cerevisiae*-based probiotic and inactivated whole yeast on vaginal candidiasis. Virulence, 8, 74–90.

Peters, B. M., Jabra-Rizk, M. A., Scheper, M. A., Leid, J. G., Costerton, J. W. & Shirtliff, M. E. 2010. Microbial interactions and differential protein expression in *Staphylococcus aureus -Candida albicans* dual-species biofilms. FEMS Immunology and Medical Microbiology, 59, 493–503.

Ribeiro, F. C., De Barros, P. P., Rossoni, R. D., Junqueira, J. C. & Jorge, A. O. 2017. Lactobacillus rhamnosus inhibits *Candida albicans* virulence factors *in vitro* and modulates immune system in *Galleria mellonella*. Journal Appllied Microbiology, 122, 201–211.

Schulenburg, H., Kurz, C. L. & Ewbank, J. J. 2004. Evolution of the innate immune system: the worm perspective. Immunological Reviews, 198, 36–58.

Singkum, P., Muangkaew, W., Suwanmanee, S., Pumeesat, P., Wongsuk, T. & Luplertlop, N. 2019. Suppression of the pathogenicity of *Candida albicans* by the quorum-sensing molecules farnesol and tryptophol. The Journal of General and Applied Microbiology, 65, 277–83.

Tobudic, S., Lassnigg, A., Kratzer, C., Graninger, W. & Presterl, E. 2010. Antifungal activity of amphotericin B, caspofungin and posaconazole on *Candida albicans* biofilms in intermediate and mature development phases. Mycoses, 53, 208–14.

Van Rijswijck, I. M. H., Dijksterhuis, J., Wolkers-Rooijackers, J. C. M., Abee, T. & Smid, E. J. 2015. Nutrient limitation leads to penetrative growth into agar and affects aroma formation in *Pichia fabianii, P. kudriavzevii* and *Saccharomyces cerevisiae*. Yeast, 32, 89–101.

Wilson, D., Naglik, J. R. & Hube, B. 2016. The missing link between *Candida albicans* hyphal morphogenesis and host cell damage. Plos Pathogens, 12.

Zhao, Y. F., Wu, J. F., Shang, D. R., Ning, J. S., Ding, H. Y. & Zhai, Y. X. 2014. Arsenic species in edible seaweeds using *in vitro* biomimetic digestion determined by High-Performance Liquid Chromatography inductively coupled plasma Mass Spectrometry. International Journal of Food Science, 2014, 1–12.

