## Supplemental table 1 for "Secondary metabolites from food-derived yeasts inhibit virulence of *Candida albicans*"

**Table supplement 1.** Yeast and bacterial strains used in the study.

| **Yeasts/ bacteria** | **Strain name** | **Accession number/ deposition number** |
|---|---|---|
| *Saccharomyces cerevisiae* | KTP | KY47139 1 |
| *Saccharomyces cerevisiae* | F1322 | Chen and Fink, 2006 |
| *Saccharomyces cerevisiae* | L8184 | Chen and Fink, 2006 |
| *Saccharomyces cerevisiae* var *boulardii* | | NCDC 363 |
| *Issatchenkia occidentalis* | ApC | KF551991 |
| *Staphylococcus aureus* | - | FRI722 |
| *Micrococcus luteus* | - | ATCC9341 |
| *Pseudomonas aeruginosa* | - | MTCC 2297 |
| *Bacillus subtilis* | - | MTCC 736 |

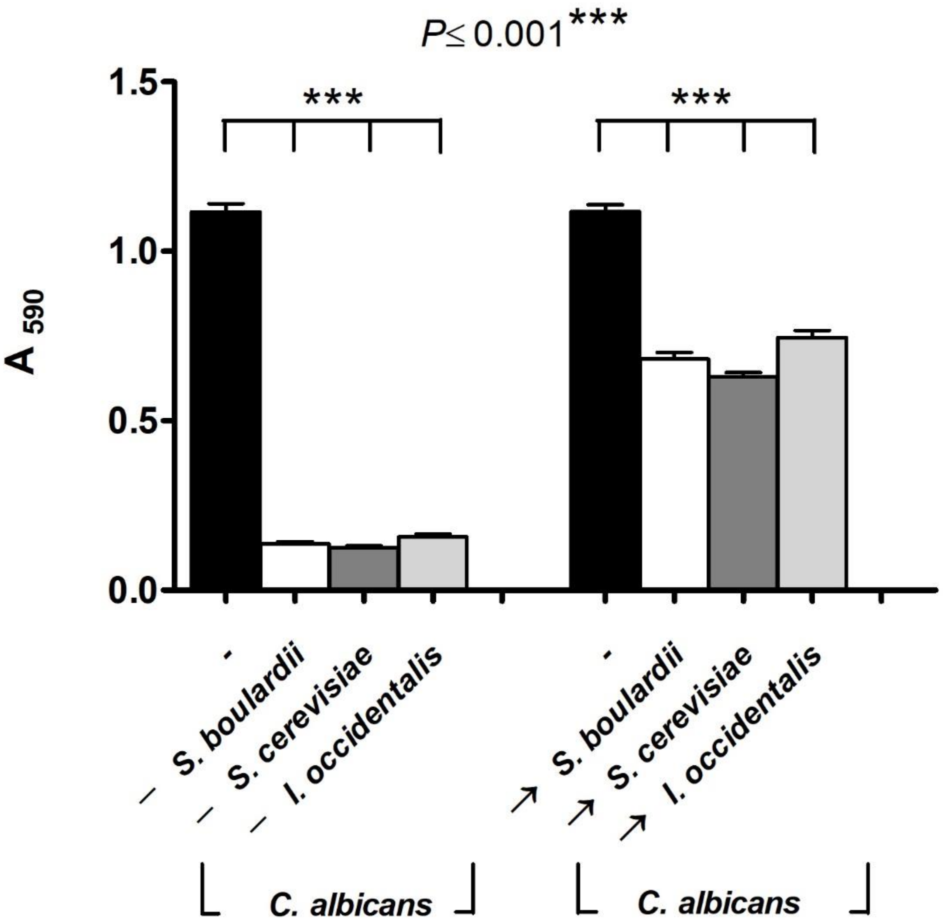


**Figure supplement 1.** The novel, food derived yeasts strains *S. cerevisiae*, *I. occidentalis* were pre-inoculation (-) and post-inoculation (→) with *C. albicans* in RPMI and/or SC media. 0.5% crystal violet was used for quantifying the adhered *C. albicans* cells on abiotic surfaces. *S. boulardii* was used as a reference condition for the purpose of comparison (n=8 assay replicates). Error bars represents means± standard deviations (SD) and all values were represented at a significance with respect to control using Tukey's *t* test.

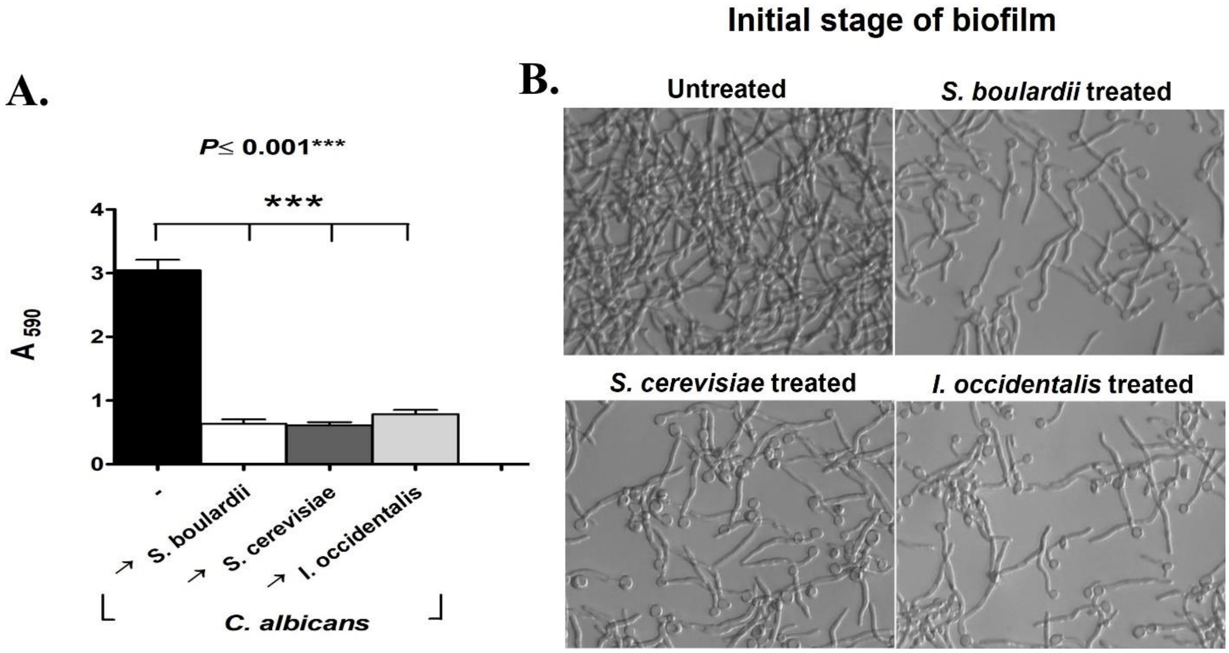


**Figure supplement 2.** (A) Effect of the beneficial yeasts on preformed *C. albicans* biofilms (up to 5-6 h old), under post-inoculation conditions. Adhesion was quantified using 0.5 % crystal violet staining after 24 h. (B) Micrographs showing *C. albicans* early stages of biofilm formation on abiotic surfaces treated with the beneficial yeasts *S. cerevisiae*, *I. occidentalis* and compared to the reference strain *S. boulardii* (n=8 assay replicates). All values represent means± standard deviations (SD), significance levels were expressed with respect to control using Tukey's *t* test.

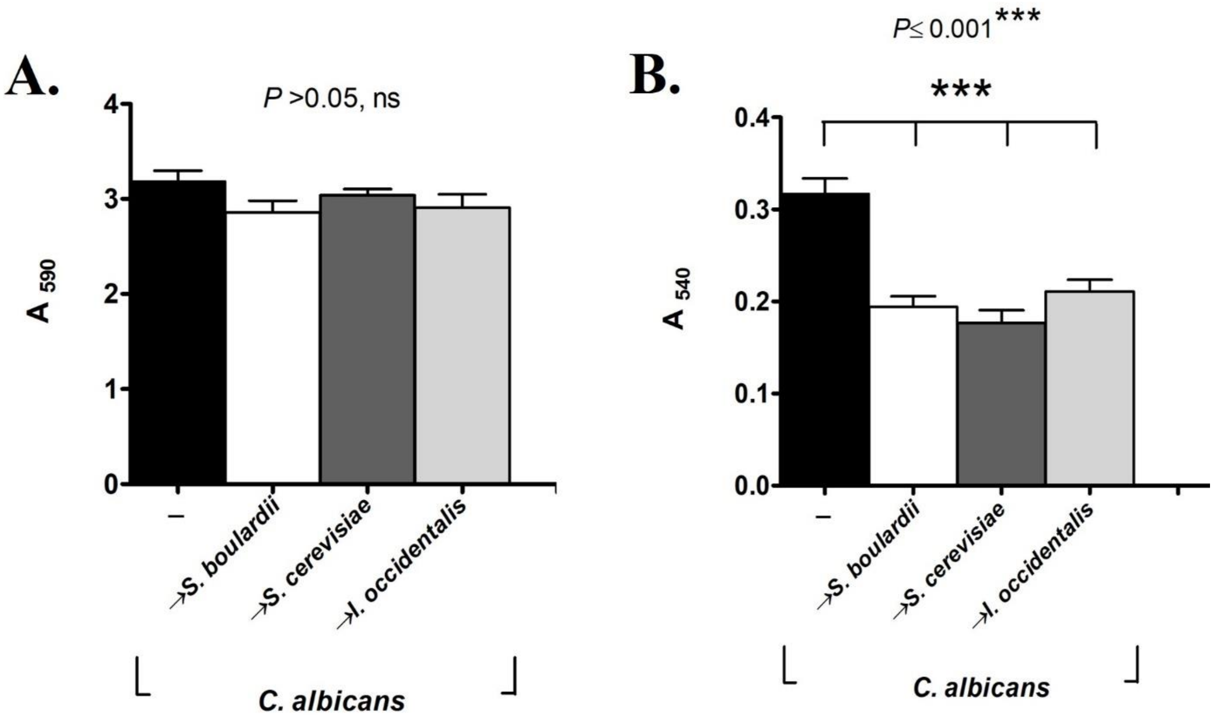


**Figure supplement 3.** (A). Exposure of the beneficial yeasts *S. cerevisiae*, *I. occidentalis* or the reference strain *S. boulardii* to 24 h old *C. albicans* biofilms, did not show any reduction in biomass but (B) exhibited a marked decrease in metabolic activity (n= 4 assay replicates).

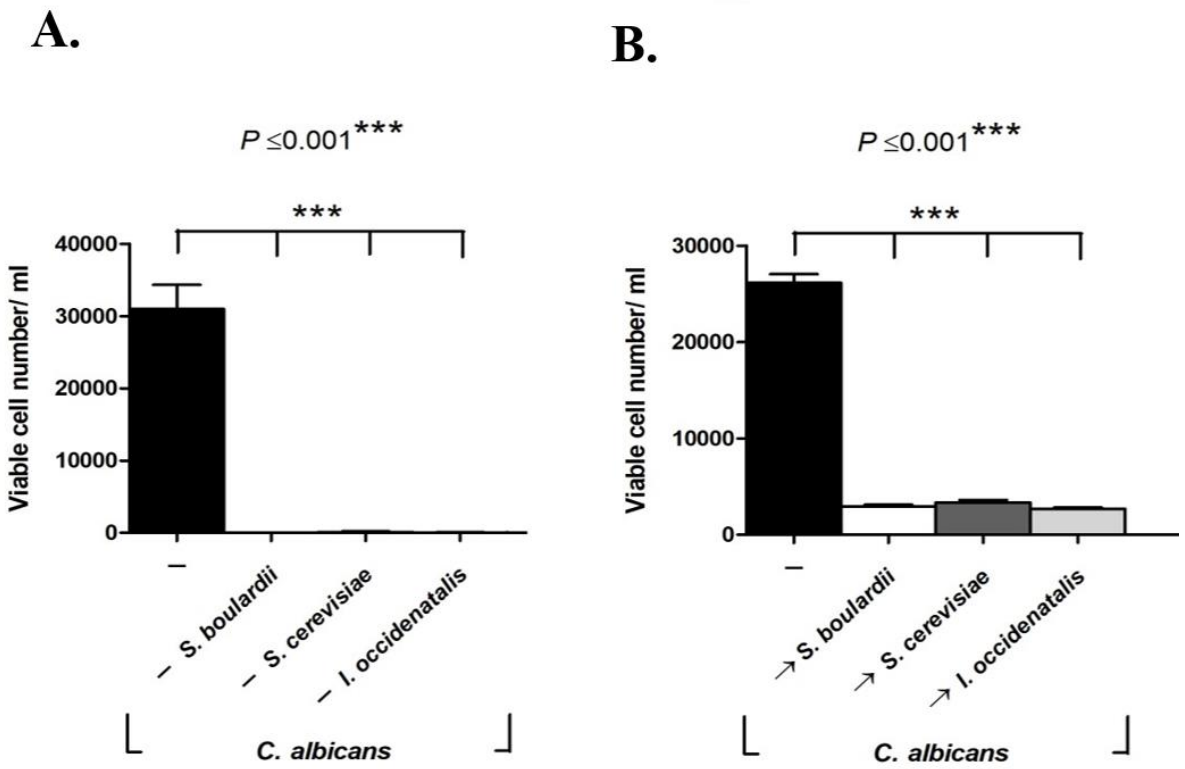


**Figure supplement 4.** Pre-inoculated (represented as -) (A.) and post-inoculated (→) (B.) beneficial yeast against *C. albicans* reduced the virulence on Ccao-2 monolayer cell line (n= 3 assay replicates).

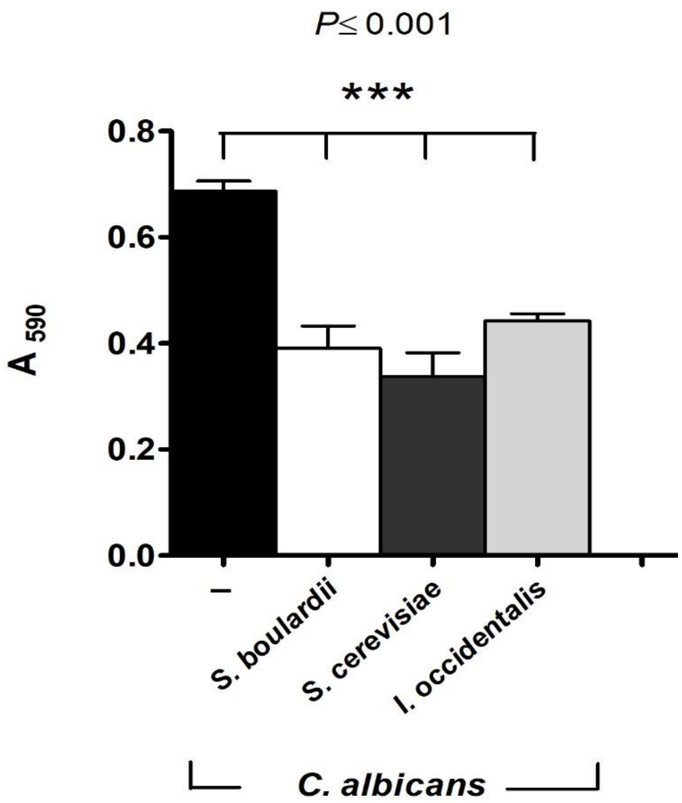


**Figure supplement 5.** Beneficial yeasts are able to inhibit adhesion of *C. albicans* in simulated bile juice even when applied 2 h after *C. albicans* (n=4 assay replicates).

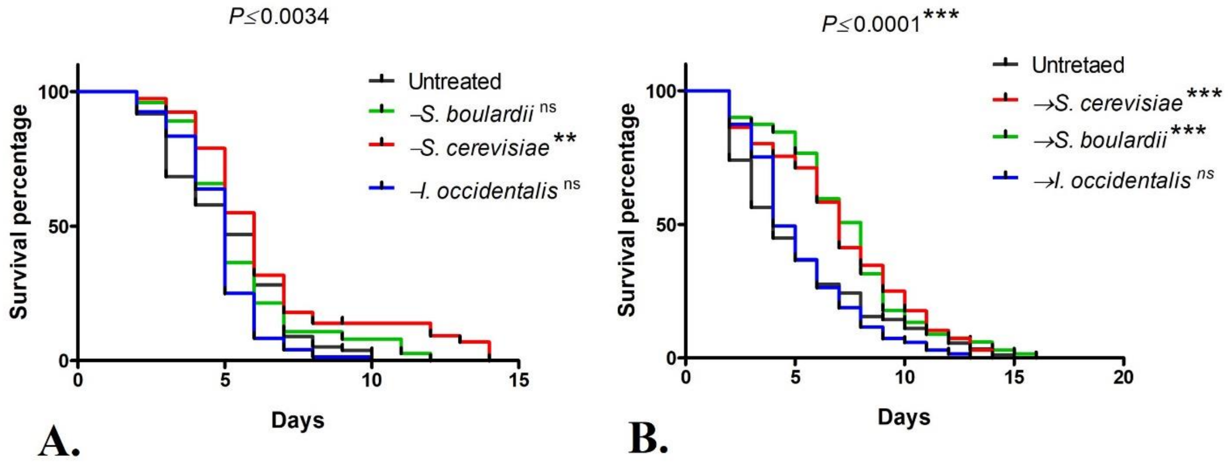


**Figure supplement 6.** (A) Lifespan of *C. elegans* infected with *C. albicans* that were pre-treated with beneficial yeasts (n=3, 108± 35 nematodes) and (B) treated after *C. albicans* infection (n=3 assay replicates, 121± 13 nematodes) on. Kaplan-Meier statistical analysis tools by Log-rank (Mantel-Cox) tests were used for the *C. elegans* survival assay. (Pre-treated worm represented as '→' and post-treated as ('−').

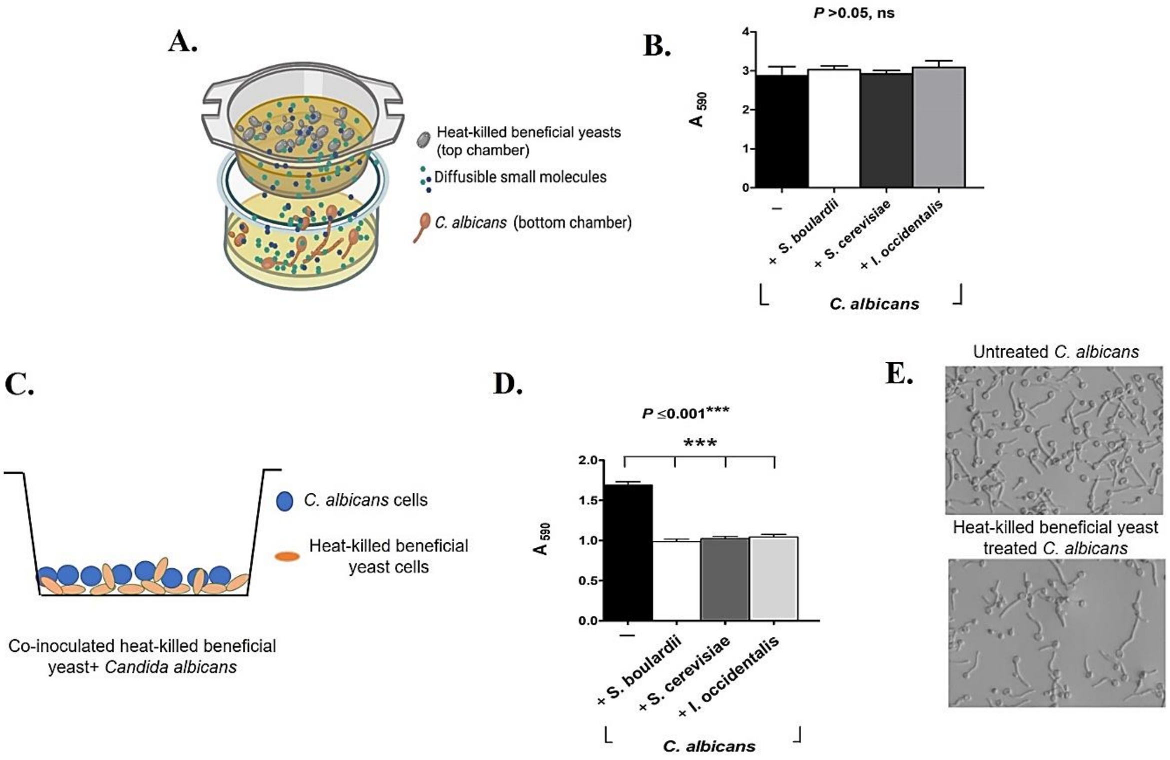


**Figure supplement 7.** Effect of heat killed beneficial yeast on *C. albicans*. (A) Heat-killed beneficial yeasts *S. cerevisiae*, *I. occidentalis* and *S. boulardii* and live *C. albicans* were inoculated to upper chamber and lower chamber of the dual-cell insert respectively (n= 4, assay replicates). (B) After incubation lower *C. albicans* inoculated chamber was washed three times with PBS and stained with 0.5 % crystal violet. (C) To understand the physical barrier effect of beneficial yeasts, (D) both heat-killed beneficial yeasts and live *C. albicans* were co-inoculated and quantified by 0.5 % crystal violet (n= 8 assay replicates). (E) An illustrative image of physical barrier effect of beneficial yeast, where heat-killed yeast reduced the adhesion of *C. albicans*, however did bot effect against filamentation.

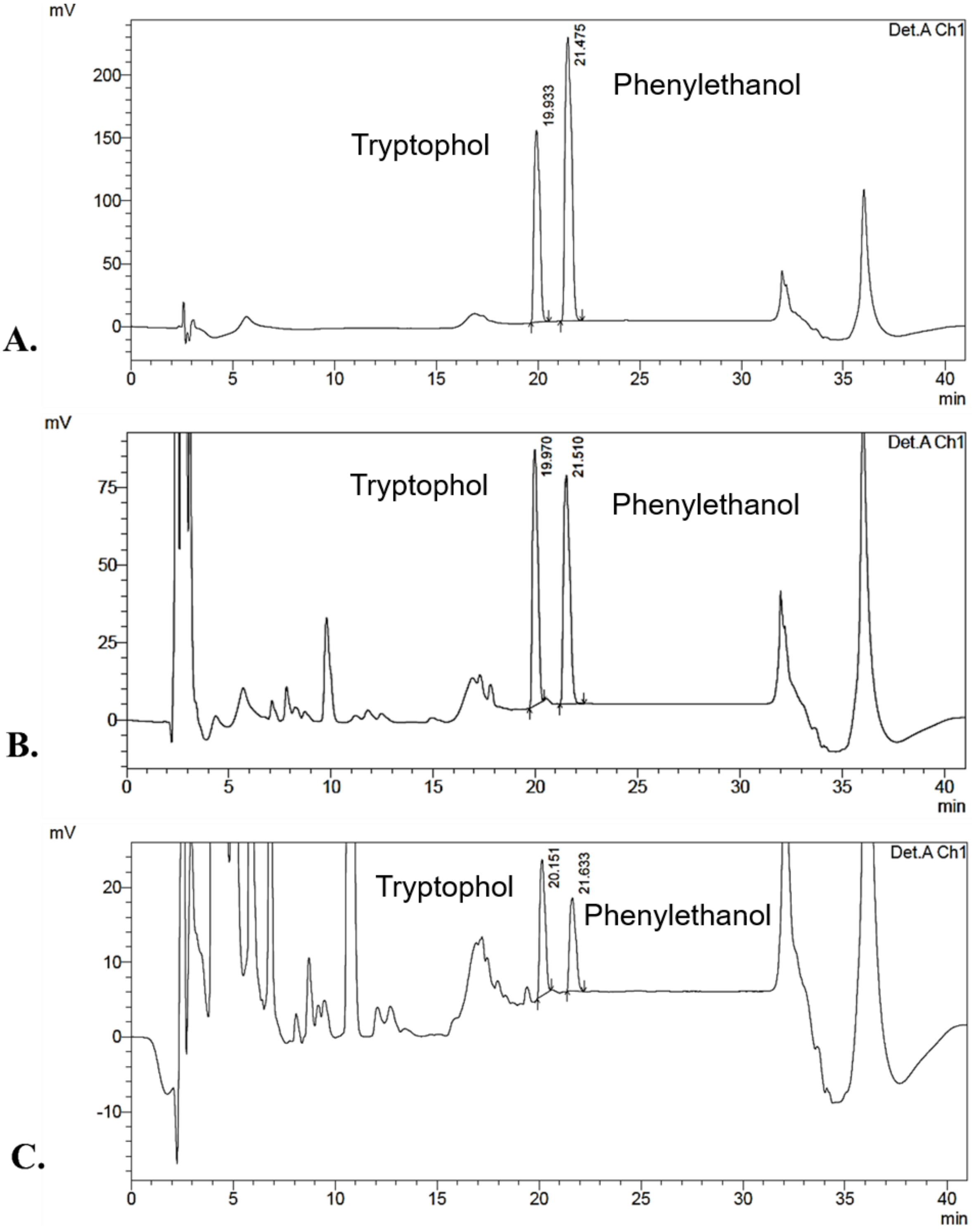


**Figure supplement 8.** High Pressure Liquid Chromatography (HPLC). (A) Representative HPLC chromatograms of phenylethanol and tryptophol of standards, (B) of beneficial yeast (*I.*

*occidentalsis*) secretome and (C) *S. cerevisiae* mutant *aro8 aro9* that is defective in phenylethanol and tryptophol production.
